# NRX-101 (D-Cycloserine + Lurasidone), a Qualified Infectious Disease Product, is Active Against Drug-Resistant Urinary Pathogens *In Vitro*

**DOI:** 10.1101/2024.01.14.575572

**Authors:** Michael T. Sapko, Michael Manyak, Riccardo Panicucci, Jonathan C. Javitt

**Affiliations:** NRx Pharmaceuticals 1201 N Market St Suite 111, Wilmington, DE 19801; George Washington University, Washington, DC; Johns Hopkins University, Baltimore, MD

**Keywords:** d-cycloserine, complicated urinary tract infections, susceptibility testing, antimicrobials, NRX-101, multidrug resistance

## Abstract

**Background:** D-Cycloserine (DCS) is a broad-spectrum antibiotic that is currently FDA-approved to treat tuberculosis (TB) disease and urinary tract infection. Despite numerous reports showing good clinical efficacy, DCS fell out of favor as a UTI treatment because of its propensity to cause side effects. NRX-101, a fixed dose combination of DCS and lurasidone, has been awarded Qualified Infectious Disease Product and Fast Track Designation by the US Food and Drug Administration and is being developed for various CNS indications because of its unique synergistic effect; each component mitigates side effects of the other.

**Methods:** In this study, we tested NRX-101 against the urinary tract pathogens *E. coli, P. aeruginosa, K. pneumoniae*, and *A. baumannii* in Mueller Hinton broth (caMHB) and artificial urine media (AUM). Several strains were multidrug resistant. Test compounds were serially diluted in broth/media. Minimum inhibitory concentration (MIC) was defined as the lowest concentration of test compound at which no bacterial growth was observed.

**Results:** DCS exhibited antibacterial efficacy against all strains tested while lurasidone did not appreciably affect the antibacterial action of DCS *in vitro*. In AUM, the MICs ranged from 128 to 512 mcg/ml for both DCS and NRX-101. In caMHB, MICs ranged from 8 to 1024 mcg/ml for NRX-101 and 32 to 512 mcg/ml for DCS alone.

**Conclusions:** Our data confirm that DCS as antibacterial activity against reference and drug-resistant urinary pathogens. Furthermore, lurasidone does not interfere with DCS’s anti-microbial action *in vitro*. These results support the clinical development of NRX-101 as a treatment for complicated urinary tract infection.

## Introduction

D-Cycloserine (DCS) is a broad-spectrum antibiotic that is currently FDA-approved to treat pulmonary or extrapulmonary tuberculosis (TB) disease (active TB) and urinary tract infection (UTI).^1^ In January 2024, a fixed dose combination of DCS and lurasidone was awarded Qualified Infectious Disease Product Designation by the US FDA for treatment of complicated Urinary Tract Infection (cUTI) and pyelonephritis. In the 1950s and 60s, DCS was used to treat various types of infections including UTI with numerous reports showing its clinical efficacy.^2-5^ Landes et al., for example, reported good clinical efficacy against a wide variety of urinary tract pathogens including *E. coli, Pseudomonas, Aerobacter aerogenes* (renamed Enterobacter aerogenes and genetically closest to *Klebsiella*), various *Proteus* strains, *Staphylococcus* and *Streptococcus*. The use of DCS as an antibiotic for the treatment of UTI fell out of favor in the 1970s. However, because the incidence of multi-drug-resistant urinary tract pathogens^6^ is outpacing the development of new antimicrobial agents^7^, several authors have begun to re-examine the use of older antibiotics such as DCS.^8,9^

D-cycloserine (DCS) inhibits two sequential enzymes in the bacterial cell wall peptidoglycan biosynthetic pathway, which forms the dipeptide D-alanyl-D-alanine (D-Ala-D-Ala).^10^ DCS targets alanine racemase to form an aromatized DCS-PLP adduct, which irreversibly blocks alanine racemase activity.^11^ DCS also competitively and reversibly inhibits D-Ala-D-Ala ligase.^10^ DCS also antagonizes a third bacterial enzyme, D-amino acid dehydrogenase.^8^

DCS is a partial agonist of the human NMDA receptor and crosses the blood-brain barrier. It is binding and activation profile that has attracted vigorous research interest in DCS as a treatment for acute suicidality in bipolar depression, post-traumatic stress disorder, and chronic pain. However, DCS action at NMDA receptors is also likely the cause of rare but potentially treatment-limiting side effects such as hallucinations.

Nonclinical and limited clinical testing conducted to date suggests the combination of DCS and lurasidone has a potential synergy in that one agent ameliorates known side effects of the other. As a combined D_2_/5HT_2A_ antagonist, lurasidone may have the ability to reverse the psychotomimetic effects of DCS, and DCS, in turn, appears to block the anxiogenic effects of lurasidone.^12,13^

NRX-101 is a fixed-dose combination of DCS and lurasidone that is being developed for various CNS indications. In the current study, we show that DCS alone and NRX-101 (DCS plus lurasidone) are active against common urinary tract bacterial pathogens, including microbes that are highly resistant to standard antimicrobials.

## Methods

### Bacterial Isolates

A total of 13 bacterial isolates were tested; 6 *Escherichia coli*, 4 *Pseudomonas aeruginosa*, 1 *Klebsiella pneumoniae* and 2 *Acinetobacter baumannii* strains. Specific strains and their antibiotic susceptibilities are listed in Appendix 1.

### Antimicrobial Susceptibility Testing

Antimicrobial susceptibility testing carried out by Charles River Laboratories (Portishead) in accordance with CLSI guidelines M07-A10. Bacterial inoculum was adjusted to the equivalent OD_600_ of 0.5 McFarland standard and added to plates at one concentration (1:1, v:v). Assays were carried out in both cation-adjusted Mueller Hinton broth (caMHB; Millipore) and artificial urine media (AUM; Biochemzone). Bacteria grown in caMHB and in AUM required 24- and 48-hours incubation time, respectively. 96-well plates with triplicate technical replicates of each condition were incubated at 37°C under aerobic conditions. To assess the minimum inhibitory concentrations (MICs), test compounds were serially diluted in broth/media to 12 dilutions. Each strain was tested for its susceptibility to d-cycloserine (DCS; Sigma-Aldrich), lurasidone (MedChemExpress), DCS + lurasidone, and one antibiotic against which the test strain is known to be susceptible. Control antibiotics gentamicin sulfate salt, colistin sulfate salt, ciprofloxacin, and tobramycin were obtained from Sigma-Aldrich. Each strain was also allowed to grow without the presence of any test compound or antibiotic. Bacterial growth was determined visually. MIC was defined as the lowest concentration (i.e., greatest dilution) of test compound at which no bacterial growth was observed. Sterility checks were performed by adding media with inoculum to each well per guidelines.

Back-plating was performed to confirm pure inoculum.

## Results

All bacterial isolates grew in CaMHB and with the exception of *A. baumannii* 19606; all bacterial isolates grew in AUM. MIC results in CaMHB and in AUM are shown in Table 1 and Table 2, respectively. DCS showed activity against all the bacterial isolates tested. Lurasidone alone had no antibacterial activity at the concentrations tested in either medium. When tested as a combination, lurasidone did not appreciably affect the antibacterial efficacy of DCS. MICs of DCS were often higher for bacterial isolates grown in AUM compared to these grown in caMHB. This effect was most apparent for the *E. coli* and *P. aeruginosa* strains tested. Reference antibiotics performed as expected.

**Table 1.** Minimum Inhibitory Concentrations of DCS and Lurasidone in Cation-adjusted Mueller Hinton Broth.

| Strain | Reference Antibiotic (µg/mL) | Lurasidone (µg/mL) | DCS (µg/mL) | DCS + Lurasidone (µg/mL) |
| --- | --- | --- | --- | --- |
| *E. coli* 35218 | 2<sup>a</sup> | >142.3 | 32 | 32 |
| *E. coli* 25922 | 1<sup>a</sup> | >142.3 | 32 | 32 |
| *E. coli* 700928 | 2-4<sup>a</sup> | >142.3 | 32 | 8 |
| *E. coli* 700336 | 2<sup>a</sup> | >142.3 | 32 | 32 |
| *E. coli* 2469 | 0.03-0.062<sup>b</sup> | >142.3 | 64-128 | 32 |
| *E. coli* Xen 16 | 1<sup>a</sup> | >142.3 | 64 | 32 |
| *P. aeruginosa* PA01 | 1<sup>c</sup> | >142.3 | 256 | 128 |
| *P. aeruginosa* 27853 | 0.5<sup>c</sup> | >142.3 | 256 | 128 |
| *P. aeruginosa* Xen 41 | 0.5<sup>c</sup> | >142.3 | 128 | 64 |
| *P. aeruginosa* BAA 3105 | 64<sup>c</sup> | >142.3 | 128 | 128 |
| *K. pneumoniae* 1705 | 1<sup>a</sup> | >142.3 | 128 | 128 |
| *A. baumannii* 19606 | 32<sup>a</sup> | No growth | 512 | 256 |
| *A. baumannii* 1605 | 64<sup>d</sup> | >142.3 | 512 | 1024 |
| a, Gentamicin; b, Colistin; c, Tobramycin; d, Ciprofloxacin |  |  |  |  |

**Table 2.** Minimum Inhibitory Concentrations of DCS and Lurasidone in Artificial Urine Media.

| Strain | Reference Antibiotic (µg/mL) | Lurasidone (µg/mL) | DCS (µg/mL) | DCS + Lurasidone (µg/mL) |
| --- | --- | --- | --- | --- |
| *E. coli* 35218 | 32<sup>a</sup> | >142.3 | 256 | 128 |
| *E. coli* 25922 | 256<sup>a</sup> | >142.3 | 256 | 256 |
| *E. coli* 700928 | 32<sup>a</sup> | >142.3 | 256 | 256 |
| *E. coli* 700336 | 128<sup>a</sup> | >142.3 | 128 | 256 |
| *E. coli* 2469 | 1<sup>b</sup> | >142.3 | >256 | >256 |
| *E. coli* Xen 16 | >256<sup>a</sup> | >142.3 | 256 | 256 |
| *P. aeruginosa* PA01 | 32<sup>c</sup> | >142.3 | 512 | 512 |
| *P. aeruginosa* 27853 | 32<sup>c</sup> | >142.3 | 512 | 512 |
| *P. aeruginosa* Xen 41 | 32<sup>c</sup> | >142.3 | 512 | 128 |
| *P. aeruginosa* BAA 3105 | 16<sup>c</sup> | >142.3 | 128 | 128-256 |
| *K. pneumoniae* 1705 | >256<sup>a</sup> | >142.3 | 128 | 256 |
| *A. baumannii* 19606 | No growth | No growth | No growth | No growth |
| *A. baumannii* 1605 | 32<sup>d</sup> | >142.3 | 512 | 256 |

a, Gentamicin; b, Colistin; c, Tobramycin; d, Ciprofloxacin

## Discussion

Based on the above results, the US Food and Drug Administration has awarded Qualified Infectious Disease Product and Fast Track Designation to NRX-101, a fixed dose combination of DCS and lurasidone for the treatment of complicated Urinary Tract Infection (cUTI) and pyelonephritis. DCS and DCS + lurasidone (NRX-101) showed antibacterial activity against various strains of *E. coli, P. aeruginosa, K. pneumoniae*, and *A. baumannii* in both caMHB and in AUM. In AUM, the MICs ranged from 128 to 512 mcg/ml for both DCS and NRX-101. In caMHB, MICs ranged from 8 to 1024 mcg/ml for NRX-101 and 32 to 512 for DCS alone. Kugathasan et al. tested DCS against 500 urinary pathogens and reported a range of MICs from 8 to 128 mcg/ml in Mueller-Hinton broth.^9^ That group reported the same range for the subset of 411 E. coli isolates within the larger test group. Our range of MICs against *E*.*coli* isolates 32 to 128 mcg/ml. While our group and Kugathasan’s groups both show the antibacterial efficacy of DCS, we observed lower potency of the antibiotic *in vitro*. One possible explanation is that in the large assortment of urinary pathogens Kugathasan tested, certain strains not included in our study were particularly susceptible to DCS.

Our results are congruent with those of Kugathasan et al.^14^ who demonstrated a similar range of MICs against 182 trimethoprim and 24 third-generation cephalosporin-resistant isolates. This is to be expected since DCS exerts its antibacterial effect by blocking various enzymes in the bacterial cell wall peptidoglycan biosynthetic pathway (alanine racemase activity, D-Ala-D-Ala ligase, and D-amino acid dehydrogenase). This mechanism is distinct from that of trimethoprim, which blocks the reduction of dihydrofolate to tetrahydrofolate. Similarly, the DCS mechanism is distinct from the beta-lactam action of third-generation cephalosporins against peptidoglycan transpeptidase.^8^ This distinct mechanism of action is relevant in light of the high rate of antimicrobial resistance among urinary tract pathogen isolates.

Because DCS competes with d-alanine in the peptidoglycan biosynthetic pathway to exert its antibacterial affect, various groups have suggested that DCS be tested in alanine-free media^9,14,15^ because the presence of alanine leads to “falsely elevated MICs.”^9^ Kugathasan et al. reported an MIC as low as 0.12 mcg/ml for DCS against urinary pathogenic isolates in an alanine-free, Minimal Salts medium.^9^ Indeed, the presence of alanine in culture media may explain why authors in the 1950s and 60s reported greater clinical efficacy with DCS than one would expect from corresponding MICs determined from the patient’s bacterial isolates.^3^ However, we suggest that alanine-free media provides an estimate of MIC of DCS under ideal conditions, but may not reflect the MIC under real world conditions. Both Mueller Hinton broth, derived from beef extract, and artificial urine medium used in our study contain alanine^16^ as does human urine.^17,18^

DCS is excreted mostly unchanged in the urine.^1^ Urine levels of DCS are likely to exceed the MICs for urinary pathogens reported here and elsewhere. Welch at al. reported that a single oral dose of 1,000 mg of DCS caused DCS levels in the urine to peak at approximately 200 mcg/ml at within 8 hours.^19^ From the same study, 500 mg doses of DCS given every 6 hours achieved a peak urine concentration of 800 mcg/ml at 72 hours.^19^ Fairbrother and Garrett reported similar peak urine DCS levels after 250 mg of DCS was administered orally every 8 hours in patients with urinary tract infections.^2^ We are currently performing a pharmacokinetic study of NRX-101 in plasma and urine to more precisely characterize urine levels of DCS after oral administration using modern analytical techniques.

Lurasidone did not have an effect on the anti-bacterial efficacy at any dose tested. It should be noted that the concentration of lurasidone tested *in vitro* studies, 142.3 mcg/ml, is far higher than would be expected to be found in the urine of patients taking therapeutic doses of lurasidone, which is 5.7 to 260.1 ng/ml.^20^ Thus, the inclusion of lurasidone in NRX-101 to mitigate CNS side effects from DCS is not expected to interfere with the anti-infective action of DCS in the urinary tract. The addition of lurasidone, however, is likely to offset the mild CNS effects of DCS and those effects have not been observed in studies where high-dose DCS has been used to treat bipolar depression.

## Conclusions

NRX-101, a fixed dose combination of DCS and lurasidone, is active against the urinary tract pathogens *E. coli, P. aeruginosa, K. pneumoniae*, and *A. baumannii* grown *in vitro*, including several multidrug-resistant strains. DCS exhibited antibacterial efficacy against all strains tested while lurasidone did not appreciably affect the antibacterial action of DCS *in vitro*. Urine levels of DCS are likely to exceed the MICs for urinary pathogens. These results support the clinical development of NRX-101 as a treatment for complicated urinary tract infection.

## Notes

### Competing Interest Statement

All authors have received compensation and hold equity in NRx Pharmaceuticals, Inc. The research presented was performed under contract by Charles River Laboratories, Inc.

